# NRF2-dependent epigenetic regulation can promote the hybrid epithelial/mesenchymal phenotype

**DOI:** 10.1101/2021.12.02.470615

**Authors:** Wen Jia, Mohit Kumar Jolly, Herbert Levine

**Author notes:** Correspondence (M.K.J.), (H.L.).

## Abstract

The epithelial-mesenchymal transition (EMT) is a cellular process critical for wound healing, cancer metastasis and embryonic development. Recent efforts have identified the role of hybrid epithelial/mesenchymal states, having both epithelial and mesehncymal traits, in enabling cancer metastasis and resistance to various therapies. Also, previous work has suggested that NRF2 can act as phenotypic stability factor to help stablize such hybrid states. Here, we incorporate a phenomenological epigenetic feedback effect into our previous computational model for EMT signaling. We show that this type of feedback can stabilize the hybrid state as compared to the fully mesenchymal phenotype if NRF2 can influence SNAIL at an epigenetic level, as this link makes transitions out of hybrid state more difficult. However, epigenetic regulation on other NRF2-related links do not significantly change the EMT dynamics. Finally, we considered possible cell division effects in our epigenetic regulation model, and our results indicate that the degree of epigenetic inheritance does not appear to be a critical factor for the hybrid E/M state stabilizing behavior of NRF2.

## Introduction

Epithelial-Mesenchymal Transition (EMT) is often essential for various physiological and pathological processes such as wound healing, cancer metastasis, and embryonic development (1). During EMT, cells tend to lose cell-cell adhesion and gain migration and invasion capabilities. Initially, most research assumed that EMT acted as a binary process, i.e. cells typically underwent full EMT, leading to independently-moving spindle-shaped cells. However recently, it has become clear that cells can also become stabilized in one or more hybrid epithelial/mesenchymal (E/M) states, states which exhibit a combination of epithelial and mesenchymal traits and can lead to collective cell migration and enhanced tumorigenesis (2–4).

Nuclear factor E2-related factor 2(NFE2L2, i.e. NRF2) is a transcription factor which has been shown to play an important role in preventing cells from completing full EMT during injury-induced wound healing (5). Previous mathematical models and knockdown experiments have proven that NRF2 can act as a “phenotypic stability factor” (PSF) which enables cells to more readily access a hybrid E/M state (6). In these mathematical models, NRF2 interacts with the core EMT circuit via three regulatory links: 1) NRF2 inhibits Snail (7); 2) E-cadherin inhibits the nuclear accumulation and transcriptional activity of NRF2 (8); 3) miR-200 promotes NRF2 by inhibiting Keap1 (9). All these interactions involving NRF2 act at a transcriptional or post-transcriptional level, but do not directly invoke epigenetic mechanisms such as DNA methylation or histone modification. These additional mechanisms have been shown to govern the extent and reversibility of cell-state transitions among the epithelial, mesenchymal and hybrid E/M phenotypes (10–13). Thus, the goal of this work is to ask how the aforementioned role of NRF2 can be modulated via epigenetics.

We integrated a mathematical model of coupled NRF2-EMT dynamics with a previously established phenomelogical epigenetic regulation framework (14), and studied how epigenetic regulation could affect the stability of the hybrid E/M state. Specifically, we determined the effects of adding epigenetics-based regulatory terms individually in all the three abovementioned NRF2 related-links. These terms dynamically modulate regulatory thresholds based on corresponding transcriptional activity (see Methods section). We found that incorporating epigenetic feedback affecting the inhibition of SNAIL, in other words, reducing the threshold for inhibition of SNAIL as a dynamic function of NRF2 levels, can significantly stablize the epithelial and hybrid E/M states. This effect was validated by a population dynamics analysis, and was not present for the other two links involving NRF2. Finally, we investigated the role of cell division in our model (15), and our results suggest that epigenetic fluctuations due to cell division do not appear to play an important role in altering this stabilizing behavior of NRF2.

## Results

### Epigenetic feedback on inhibition of Snail by NRF2 may stablize hybrid E/M state

In a previous study, we incorporated a phenomenological treatement of epigenetic regulation into a microRNA-based model for a core EMT circuit consist of two interconnected mutually inhibitory loops between miR-34/SNAIL and miR-200/ZEB (16). We also analyzed additional effects that occur when the additional transcription factor GRHL2 is linked to the core model (13,17) Here, we focus on an extension which adds to the core EMT model an explicit NRF2 module (**Fig 1A**). Based on experimental data, the model assumes that NRF2 inhibits Snail, while E-cad and Keap1 inhibit NRF2. E-cad and ZEB are mutually inhibitory, and miR-200 inhibits Keap1. In this model, therefore, there are three links related to NRF2, i.e. NRF2-SNAIL, NRF2-E-cad and NRF2-Keap1.

**Figure 1.**
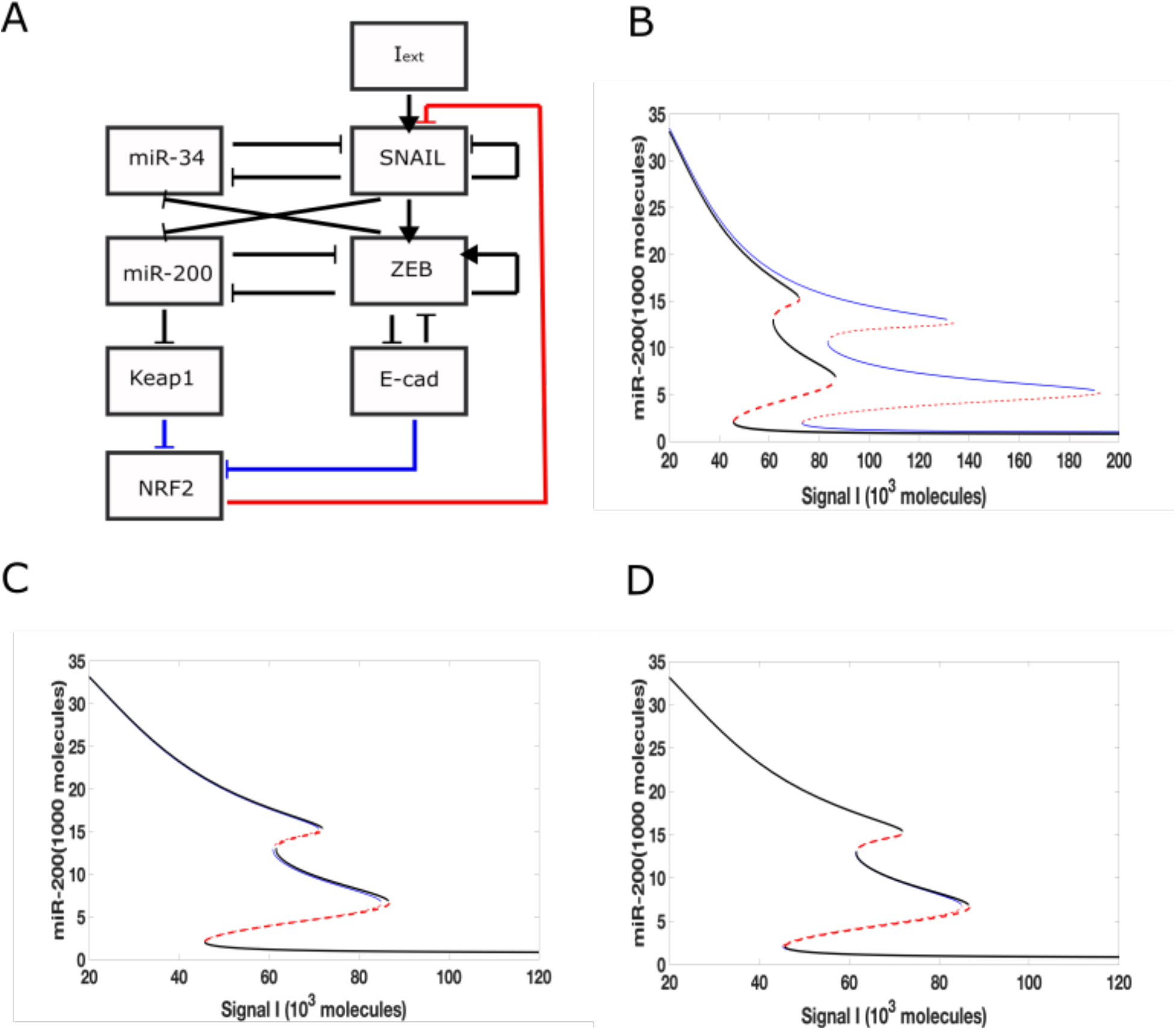
Epigenetic feedback-mediated dynamics of NRF2 coupled to core EMT circuitry. (A) A regulatory network for EMT that consists of two microRNA-TF mutually inhibitory circuits : miR-34/SNAIL and miR-200/ZEB. Signal I represents external EMT-inducing signals such as HGF, NF-κB, Wnt, TGF-β and/or HIF1α. NRF2 module is added to the core networks by three distinct links. Blue links represent the inhibition from Keap1 and E-cad on NRF2 respectively, and the red link represents the inhibition on SNAIL from NRF2.(B-D) Bifurcation diagrams of miR-200 levels for the network shown in Fig 1A, with I as the bifurcation parameter. Solid lines represent stable states, dashed lines represent unstable states. Black lines correspond to the circuit without epigenetic regulation, and blue lines correspond to a circuit including epigenetic feedback. In B), the epigenetic feedback is on the inhibition of SNAIL by NRF2. In C), the epigenetic feedback is on the inhibitory link from E-cadherin to NRF2. In D), the epigenetic feedback is on the inhibtion of NRF2 by Keap1.

We added epigenetic feedback individually to each link, and varied its strength in order to determine the range of possible effects. In each case, we first quantified these effects by deriving a new steady-state bifurcation diagram displaying how the allowed EMT states vary with an external inducing signal. In one of the three resultant bifurcation diagrams (**Fig 1B**), where epigenetic feedback regulates the inhibition acting from NRF2 on SNAIL, the range of hybrid E/M state existence is greatly increased (miR-200 > 15,000 molecules: epithelial state; miR-200 levels < 5,000 molecules: mesenchymal state; 5,000 < miR-200 <15,000 molecules: hybrid E/M state). Importantly, this increase is due to its “rightward” extension, which means even if the external EMT-inducing signal is very strong, the cell is still able to maintain its hybrid state and not automatically switch to a fully mesenchymal phenotype. In contrast, when epigenetic feedback is added to either of the two links which inhibit NRF2, the bifurcation diagrams are relatively unchanged as compared to the case when there is no epigenetic feedback (**Fig 1C-D**). Based on these results, we focus below on the inhibition link from NRF2 to SNAIL.

### Dynamic and population analysis indicate the stabilization of hybrid E/M state

Based on results from the bifurcation analysis, we expect that including epigenetic feedback on the inhibition of SNAIL by NRF2 will make a significant difference in dynamical simulations which include noise, thereby allowing cell-state transitions to spontaneously occur (18,19). Here, we start with a cell in the epithelial state, and the initial signal is set to be at the median of the tristable region (i.e. the co-existence region of E, M and hybrid E/M states) for the control (no epigenetic feedback) case. Specifically, the tristable signal region only ranges from 65,000-74,000 molecules, so we choose the initial signal to be 69,000 molecules in order to make it possible for the cell to switch among all three states. In a dynamical simulation without any epigenetic feedback (**Fig 2A**), most of the time the cell stays in a mesenchymal state (high ZEB, low miR-200). Intsead, if we include the epigenetic effect (**Fig 2B**), the cell spends increased amounts of time in either a hybrid state (medium ZEB, medium miR-200) or an epithelial state (low ZEB, high miR-200), thus reflecting a significant change in the mean residence time (MRT) of the three phenotypes. These results are reminiscent of changes seen in MRT for these phenotypes upon including other “phenotypic stability factors” similar to NRF2 (20).

**Figure 2.**
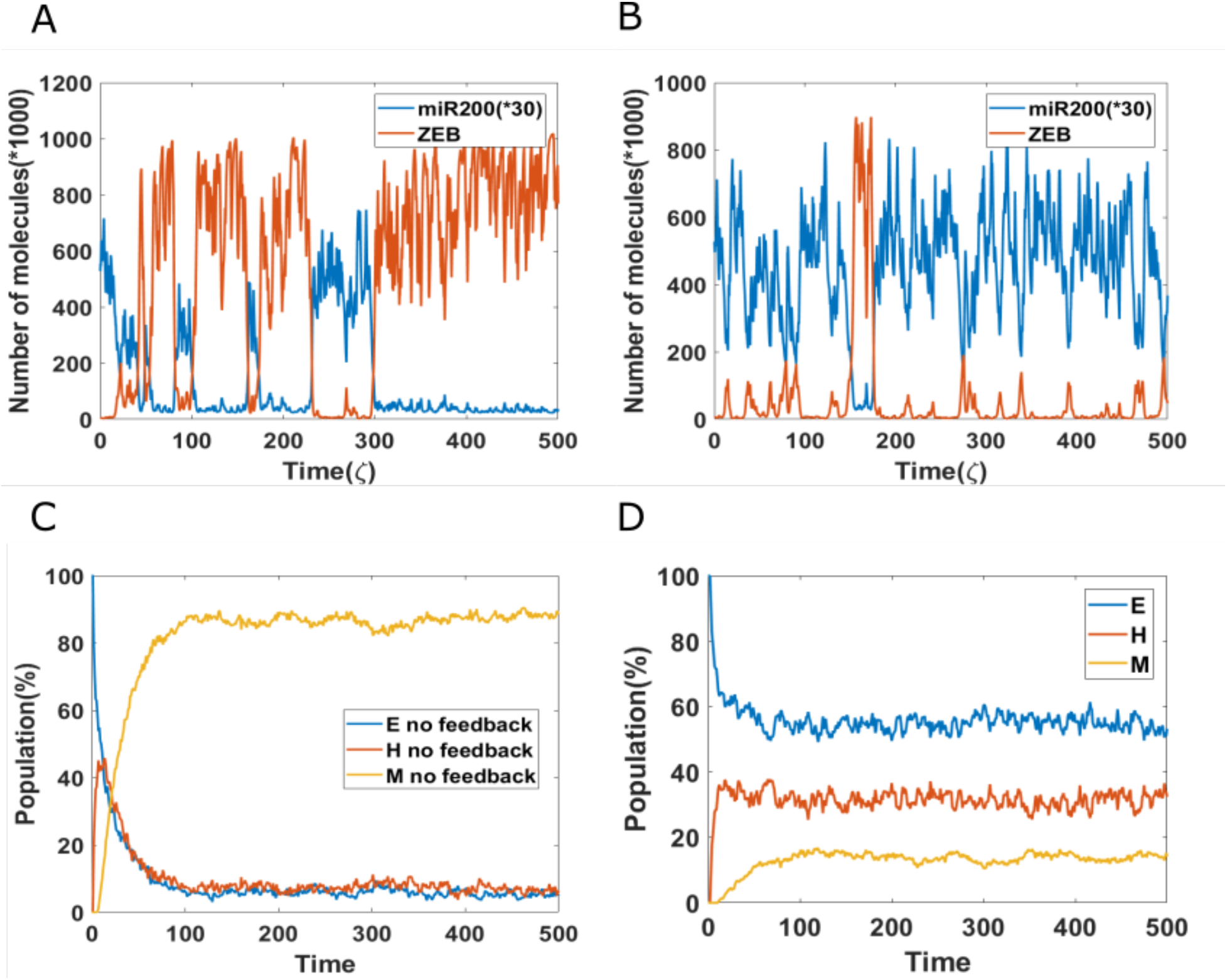
Population dynamics of NRF2 mediated epigenetic feedback. (A) A sample dynamic plot without epigenetic feedback. (B) A sample dynamic plot with feedback on the inhibition of SNAIL by NRF2. (C) Simulations showing the population change as a function of time. The percentage is calculated based on 1000 independent simulations. There is no epigenetic regulation. (D) Same as C) but now including epigenetic feedback on the inhibitory link from NRF2 to SNAIL.

These trajectory results have direct implications for the population dynamics. To see this, we performed population simulations with and without epigenetic feedback. Here, each cell had their initial phenotype as epithelial, and the initial signal is again set equal to the median of the tristable region for the bifurcation diagram obtained without any epigenetic feedback. In case of no epigenetic feedback, the population reaches a steady-state distribution of 4.5% epithelial, 7.8% hybrid and 87.7% mesenchymal (**Fig 2C**). In comparision, when epigenetic feedback is added on the inhibition of SNAIL by NRF2, the population distribution becomes 52.4% epithelial, 31.4% hybrid and 16.2% mesenchymal (**Fig 2D**). This dramatic increase in the population percentage of cells in the epithelial or hybrid states is consistent with the individual cell dynamics (**Fig 2A-B**).

### Epigenetic feedback by NRF2 stablizes hybrid state when competing with mesenchymal state

So far, we have investigated the stabilization of hybrid and epithelial states. Following that, we now study the competition between hybrid state and mesenchymal state. We chose the mean of signal to be 140K, which is mainly in the {E/M, M} region for models both with or without epigenetic feedback cases, and which lies close to the edge of the epithelial state (**Fig 3A**). We implemented a population analysis for 100 cells. Without epigenetic feedback, the percentage of hybrid E/M cells continues decreasing when the percentage of mesenchymal is increasing, and the population reaches a distribution consisting of 5%E, 10% E/M, 85% M at the end of our simulation (**Fig 3B**). However, with the epigenetic modification, the population maintains a high level of hybrid, and still has 84% E/M in the end, while the E and M stabilize around 8% respectively (**Fig 3C**). Given the huge difference revealed by this analysis, we suggests that epigenetic feedback by NRF2 may help cells maintain a hybrid state, when mainly competing with mesenchymal fates.

**Figure 3.**
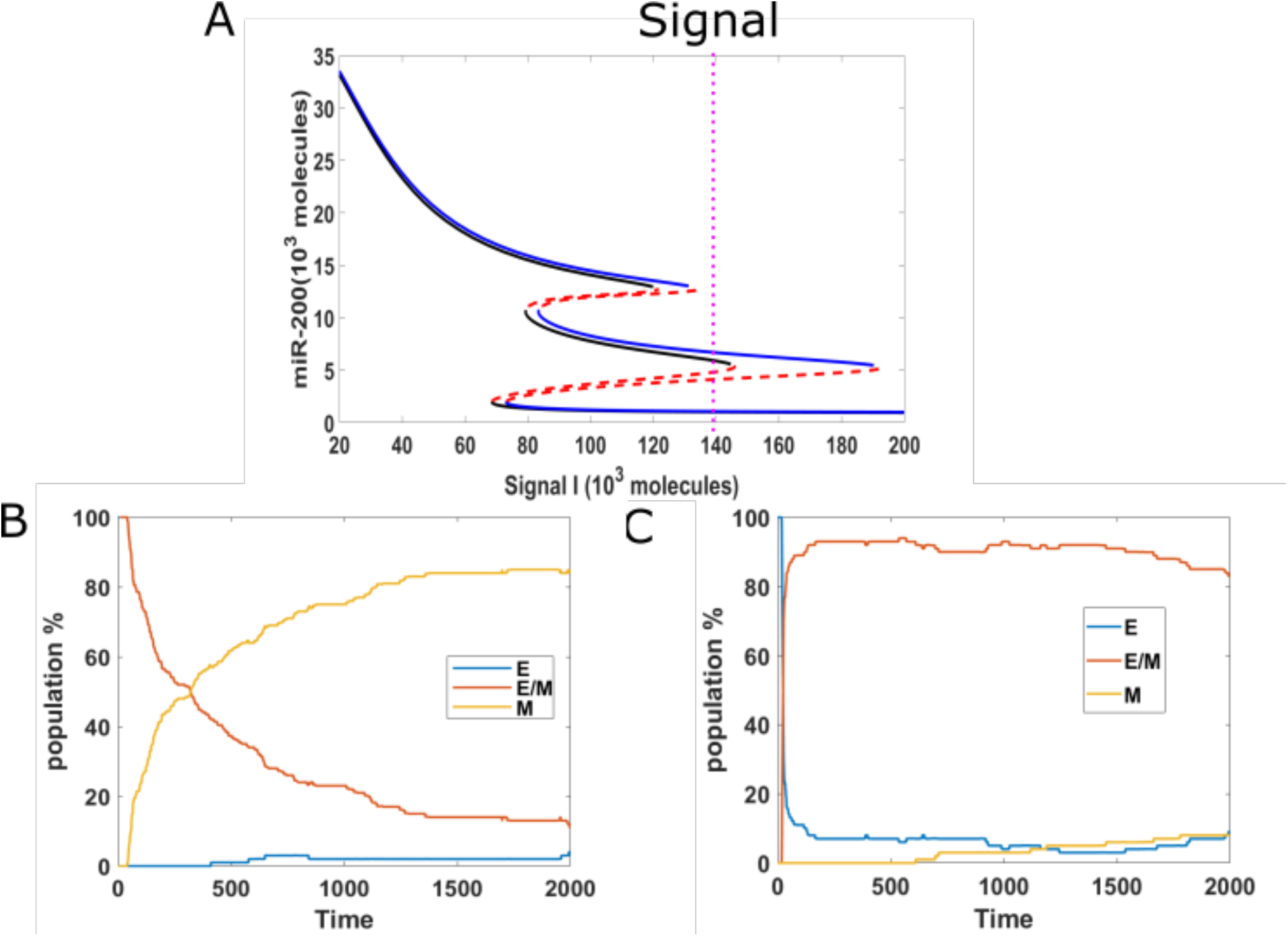
(A) Bifurcation results(black lines represent no feedback, blue lines represent feedback on NRF2’s inhibition on SNAIL, and the mean of signal is fixed at 140K (purple dotted line). (B) Population analysis without epigenetic feedback for 100 cells. (C) Population analysis with epigenetic feedback for 100 cells.

### Epigenetic feedback by NRF2 helps cells maintain epithelial state when competing with hybrid state

Next, we investigated the competition between epithelial and hybrid states. In order to increase the range of the tristable region, we added a GRHL2 module to our baseline model (**Fig 4A-B**). We then treat cells with a relatively high external signal (I=170K, green dotted line, {H,M} region for epigenetic model, {M} region for the non-epigenetic model) for different time durations, and subsequently reduce the signal to a lower level (I=102K, purple dotted line, {E,H,M} region for both the epigenetic and non-epigenetic cases). When the signal is high for only a short time period in circuits with or without epigenetic feedback, the cells typically remain in their initial epithelial state (**Fig 4C(1), 4C(3)**), though cell may occasionally transit to hybrid state in cases without epigenetic feedback. The population results indicate a final stable distribution of 30% hybrid E/M, 70% E when without epigenetic feedback (**Fig 4C(2)**), and fully epithelial states with epigenetic feedback. When the duration of the high signal is extended in the model without epigenetic influence, the cells can reach hybrid state and stay for a certain time period (**Fig 4D(1), Fig 4D(3)**). In some samples with epigenetic feedback, cell still stay in the epithelial state (**Fig 4D(3)**). Over a long time period, the population analysis shows that in both cases the percentage of hybrid state is slowly decreasing while the percentage of epithelial state is increasing (**Fig 4D(2), Fig 4D(4)**). Meanwhile, the increase in percentage of cells in an epithelial state is faster in the case of an epigenetic feedback, and the whole population eventually has a higher percentage of epithelial cells, as compared with the no epigenetic feedback group. This comparison also indicates that the system requires a longer time to re-organize its population when epigenetic feedback plays a role. Finally, if we keep increasing the treatment duration of high signal in the baseline case, a cell can complete the full epithelial-mesenchymal transition, and maintain a mesenchymal state after signal reduction (**Fig 4E(1)**); the whole population becomes fully mesenchymal in this case as the noise level is insufficient to casue transitions. However, the population can still maintain hybrid state cells when adding epigenetic feedback (**Fig 4E(3)**), and the distribution is similar to that of **Fig 4D(4)**. These results reveals that when epigenetic feedback regulates the inhibition from NRF2 to SNAIL, the epithelial state becomes more stable when it competes with hybrid states, and also the epigenetic feedback increases the re-organizing time needed for our circuit. We again see the relative suppression of more mesenchymal states in the prersence of epigenetic effects.

**Figure 4.**
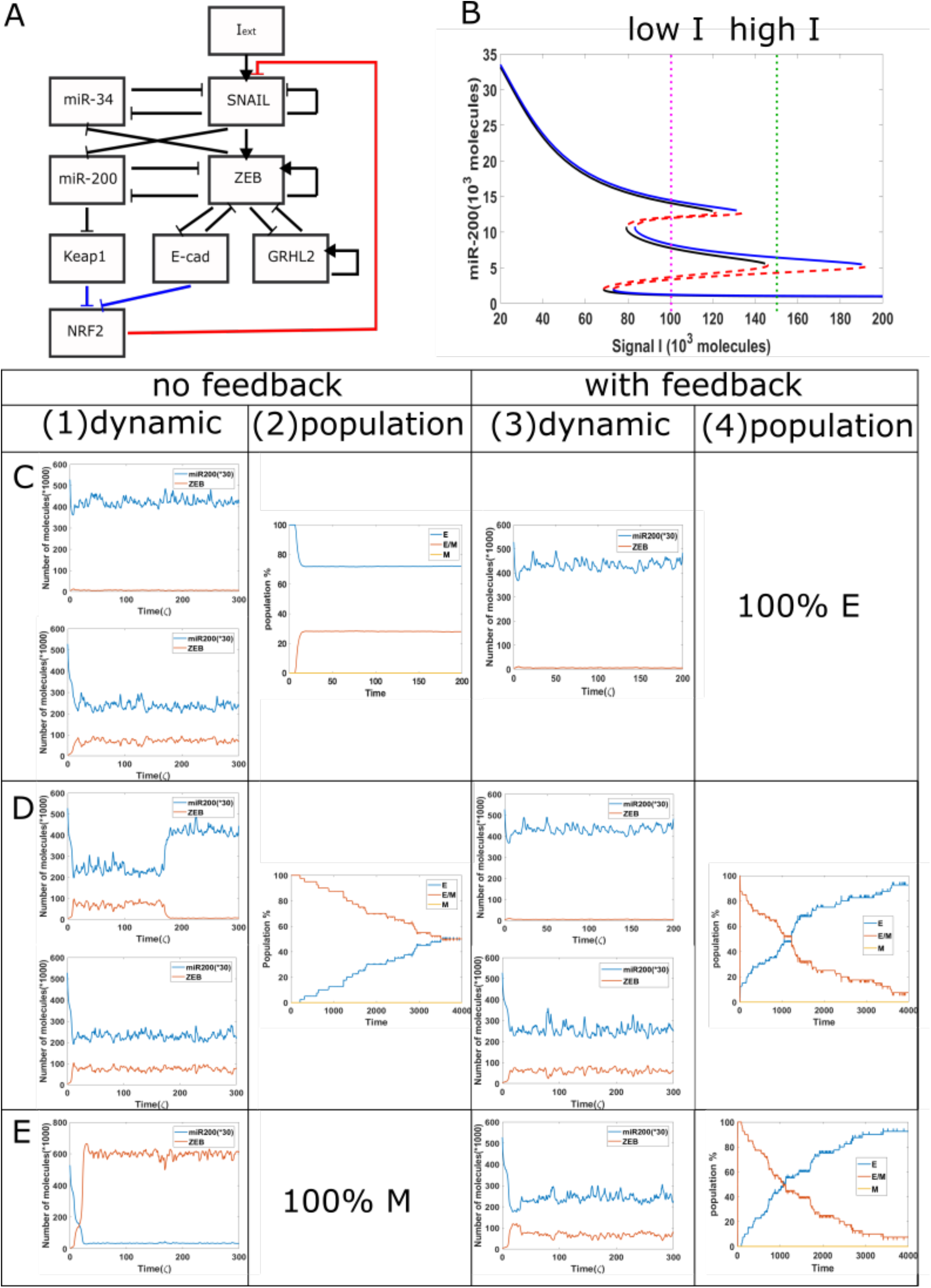
Population dynamics for the circuit including GRHL2 with/without epigenetic feedback. (A) New circuit with GRHL2. (B) Same as Fig. 3A, purple dotted line represents low signal, and green dotted line represents high signal.(C) Starting from I=170 K molecules, we show dynamic examples and population results for models with or without epigenetic feedback obtained by reducing the signal I to 102 K molecules after short high signal durations (high signal ending time=4 ς). (D) Same as Fig. 3C with high signal ending time= 10 ς. (E) Same as Fig. 3C with high signal ending time= 30 ς.

### Epigenetic inheritance may not be precisely required for the effect of NRF2 on hybrid E/M stability

Finally, we combined a previous cell division model (14)with this epigenetic regulation model, in order to see how imprecise epigenetic memory would affect the population behavior and influence the behavior of NRF2 in terms of stabilizing hybrid E/M phenotype.

In our original cell division model, the signal I_ext acquires a noise term during cell division, due to unequal distribution of signaling molecules; thus, the daughter cells may have different cell type as compared with the parent cell. Here, we considered the model containing the aforementioned epigenetic term governing SNAIL’s inhibition by NRF2, and introduced noise into the threshold value in the Hill function which governs the inhibition from NRF2 to SNAIL, during each cell division. This stochasticity mimics the possibility that epigenetic marks may not be perfectly reproduced in daughter cells. We simulated three cases with threshold noise values 0K, 100K and 300K (intial threshold =1000K). Surprisingly, all cases gave rise to similar distributions, around 50% epithelial, 13% hybrid and 37% mesenchymal (**Fig 5A**). In other words, the failure of precise epigenetic inheritance seems to have a negligible effect on the population dynamics of EMT.

**Figure 5.**
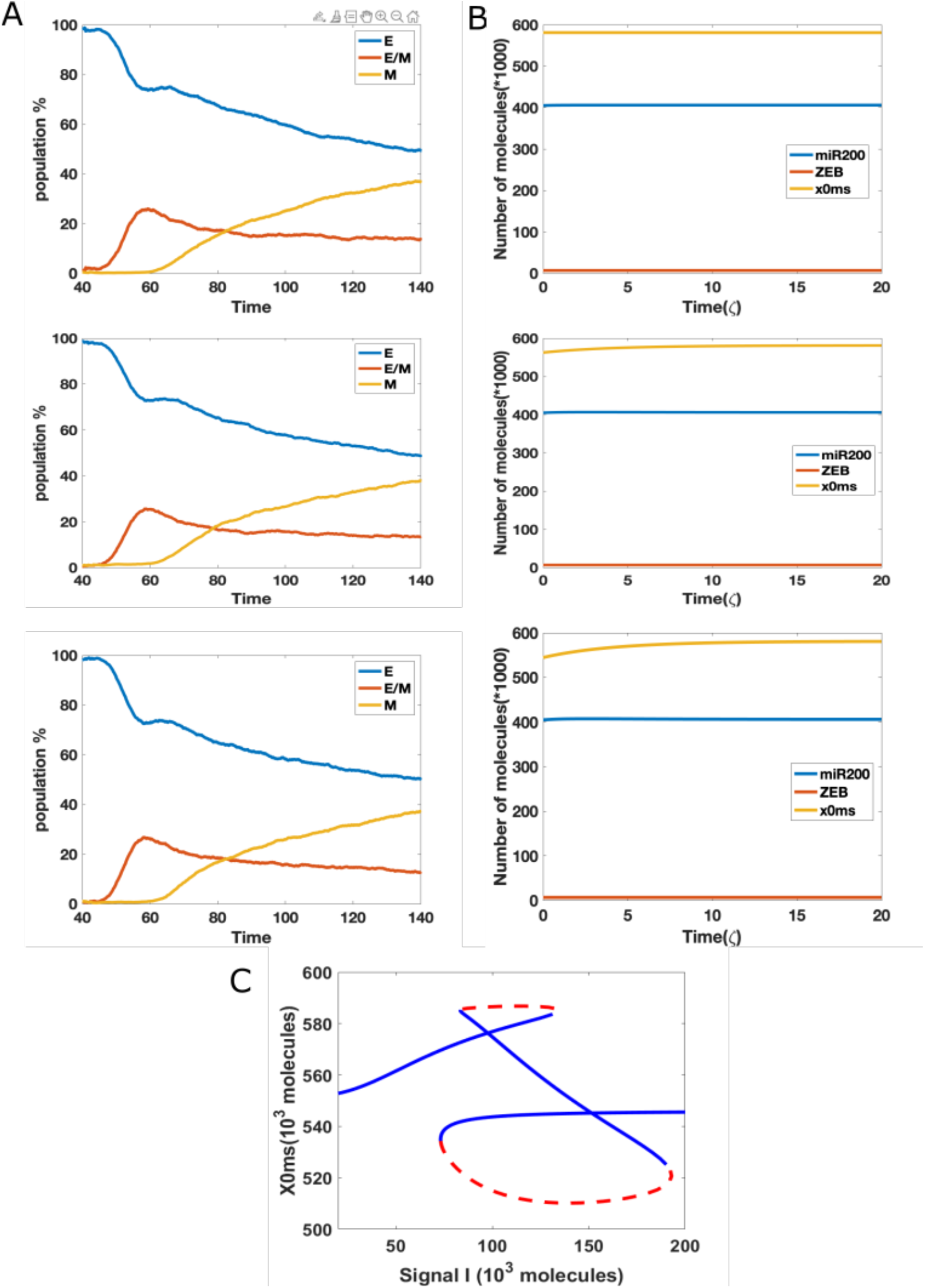
(A) Population changes as a function of time in a model including both cell division and epigenetic regulation, and where the noise of the threshold x0ms is increasing (top=0k, median=100K, bottom=300K). (B) Dynamic example for different initial x0ms values (top=581.2K, median=562.5K, bottom=544.6K; these are stable values of x0ms when I=120 from Fig. 4C). (C) Bifurcation diagram for x0ms based on inpit signal I, for the model including both epigenetic regulation on SNAIL’s inhibition from NRF2 and NRF2’s inhibition from Keap1.

In order to better understand this result, we plotted the bifurcation diagram for the threshold as a control parameter, because our simulations varied the threshold value (**Fig 5C**). We picked the value of signal I = 110K that lies in the tristable region. Then, we ran dynamical simulations to capture trends in individual cells, keping all intial parameters fixed at the epithelial state except for the threshold value. We found that only varing this threshold is not sufficient for cells to make transitions, even when the initial threshold is set to equal the value for the stable mesenchymal state (**Fig 5B**). In fact, the epigenetic dynamics is driven by the values of other variables which quickly return to their value in the epithelial state. Based on this analysis, we conclude that in this model, noise in the epigenetic state is not important in terms of determing the population structure.

## Discussion

Many computational and experimental studies have focused on a set of phenotypic stability factors such as GRHL2, OVOL2 and Np63α, which can prevent cells from undergoing the full EMT process and thus help cells maintain a hybrid epithelial/mesenchymal state (21–27). NRF2 is also one of these phenotypic stability factors, and previously constructed mechanism-based NRF2-included EMT model successfully showed that NRF2 can stablize hybrid states (6).

Here, we add addiitoanl regulatory dynamiucs due to the epigenetic effects. Our epigenetics-enhanced EMT model predicts that when there is epigenetic regulation modulating the inhibition of SNAIL by NRF2, the stabilization of hybrid states can be further enhanced. In such a case, cells can favor epithelial or hybrid E/M states, depending on the bifurcation region they lie in. Considering the epigenetic regulation, the epithelial state is more stable when competing with the hybrid state, while hybrid state is greatly favored when competing with the mesenchymal state.

We also studied the implications of our phenomelogical model of epigenetic regulation for cell division. In our simulation, the epigentic feedback regulates cells in similar timescale compared with cell division timescale. Our results indicate that error-free epigenetic inheritance may not be critical in determing EMT population balance. This does not necessarily imply that cell division is not necessary, as cell division also introduces noise in all TF and microRNA concentrations (28). In fact, recent experiments (29) seem to indicate that suppressing division does interfere with EMT, albeit without demonstrating a specific causal connection. We should note that our result may be model-dependent, and approaches in which epigenetic effects are more directly coupled to the allowed cell states (30) might give different results. We plan to further investigate this issue in future.

## Methods

The dynamics of our EMT regulatory circuit were simulated by Ordinary Differential Equations(ODES), and all the mathematical equations and corresponding parameters are shown in SI Section 1 Table 1. The simulations were implemented by Matlab.

## Acknowledgments

The work of WJ and HL was supported by the National Science Foundation sponsored Center for Theoretical Biological Physics – award PHY-2019745, and by PHY-1605817. MKJ was supported by a Ramanujan Fellowship awarded by SERB, DST, Government of India (SB/S2/RJN-049/2018).

