## Supplementary Information for "NRF2-dependent epigenetic regulation can promote the hybrid epithelial/mesenchymal phenotype"

### 1. Theoretical model for EMT

The EMT network consists of two mutually inhibiting loop: miR-200/ZEB and miR-34/SNAIL. Deterministic equations for miR-200/ZEB circuit with the external signal as SNAIL are given below [1]:

$$\dot{\mu}_{200} = g_{\mu_{200}} H^s(Z) H^s(S) - g_z H^s(Z) H^s(S) H^s(E) P_y(\mu_{200}, 6) - \gamma_\mu \mu_{200}$$
$$\dot{Z} = k_p g_z H^s(Z) H^s(S) H^s(E) P_l(\mu_{200}, 6) - \gamma_z Z$$

and those for miR-34/SNAIL circuit with I as an external signal are:

$$\dot{\mu}_{34} = g_{\mu_{34}} H^s(S) H^s(Z) - g_s H^s(X) H^s(S) H^s(I_{ext}) P_y(\mu_{34}, 2) - \gamma_{\mu_{34}} \mu_{34}$$

$$\dot{S} = k_p g_s H^s(X) H^s(S) H^s(I_{ext}) P_l(\mu_{34}, 2) - \gamma_s S$$

and for the NRF2 session:

$$\dot{K} = k_K H^s(\mu_{200}) - \gamma_K K$$

$$\dot{E} = k_E H^S(Z) - \gamma_E E$$

$$\dot{X} = k_X H^S(K) H^S(E) - \gamma_X X$$

where $g$ is the innate synthesis rate for corresponding microRNA /protein,$k_p$ is the translation rate for ZEB and SNAIL, $\gamma$ is the corresponding innate degradation rate, and $k_{K,E,X}$ is the single production rate for KEAP1, E-cadherin and NRF2. Here $H^S$ represents the shifted Hill function which is defined as:

$$H^s(B) = \frac{1 + \lambda\left(\frac{B}{B_0}\right)^{n_B}}{1 + \left(\frac{B}{B_0}\right)^{n_B}}$$

where $\lambda$ is the fold change regulated by protein B. $\lambda$>1 for activation and $\lambda$<1 for inhibition.

$P_y(\mu, n)$ describes the decrease in the level of microRNA because of the degradation of the microRNA/mRNA complex. The detailed derivation of these functions in the Supplementary Information of Lu *et al.* [2].

The external signal I that we use here can be written as the stochastic differential equation:

$$\dot{I} = \beta(I_0 - I) + \eta(t)$$

where $\eta(t)$ satisfies the condition that $< \eta(t), n(t') \geq \Gamma\delta(t - t')$. Here $I_0$ is set at 50 K molecules, $\beta$ as 0.04 hour$^{-1}$, and $\Gamma$ as 1000 (K molecules/hour)$^2$.
The initial value of I is fixed to lie at the middle of the tristable region {E, E/M, M}.

For the analysis shown in Fig 5, we used the same cell division model described previously [3], where

$$I_{sig}^{daughter} = I_{sig}^{parent} + N(0,1)\eta$$

and

$$B_{0_{sig}}^{daughter} = B_{0_{sig}}^{parent} + N(0,1)\eta.$$

**2. Epigenetic feedback regulation term**

In the EMT model, we tested epigenetic feedback through three different pathways. The dynamic equation of epigenetic feedback on NRF2's inhibition on SNIAL is:

$$\dot{X}_{m_S}^0 = \frac{X_{m_S}^0(0) - X_{m_S}^0 - \alpha X}{\zeta}$$

The other two epigenetic regulation pathway are modeled by similar method.
where $\zeta$ is a timescale factor and chosen to be 100 (hours). $\alpha$ represents the strength of epigenetic feedback. Larger $\alpha$ corresponds to stronger epigenetic feedback. $\alpha$ has an upper bound (usually between 0.01-0.3) because of the restriction that the numbers of all molecules must be positive.

**3.Parameters for the EMT model**

**Table SI 1. List of parameters used in shifted Hill functions**

| Description | Fold change | Value | # of binding sites | Value | Threshold | Value (K molecules) |
|---|---|---|---|---|---|---|
| **Inhibition on miR-200 by ZEB** | $\lambda_{Z,\mu_{200}}$ | 0.1 | $n_{Z,\mu_{200}}$ | 3 | $Z^0_{\mu_{200}}$ | 220 |
| **Inhibition on miR-200 by SNAIL** | $\lambda_{S,\mu_{200}}$ | 0.1 | $n_{S,\mu_{200}}$ | 2 | $S^0_{\mu_{200}}$ | 180 |
| **Self-activation of ZEB** | $\lambda_{Z,m_z}$ | 7.5 | $n_{Z,m_z}$ | 2 | $Z^0_{m_z}$ | 25 |
| **Activation on ZEB by SNAIL** | $\lambda_{S,m_z}$ | 10.0 | $n_{S,m_z}$ | 2 | $S^0_{m_z}$ | 180 |
| **Inhibition on miR-34 by SNAIL** | $\lambda_{S,\mu_{34}}$ | 0.1 | $n_{S,\mu_{34}}$ | 1 | $S^0_{\mu_{34}}$ | 300 |
| **Inhibition on miR-34 by ZEB** | $\lambda_{Z,\mu_{34}}$ | 0.2 | $n_{Z,\mu_{34}}$ | 2 | $Z^0_{\mu_{34}}$ | 600 |
| **Self-inhibition of SNAIL** | $\lambda_{S,m_s}$ | 0.1 | $n_{S,m_s}$ | 1 | $S^0_{m_s}$ | 200 |
| **Activation on SNAIL by external signal I** | $\lambda_{I,m_s}$ | 10 | $n_{I,m_s}$ | 2 | $I^0_{m_s}$ | 50 |
| **Inhibition on SNAIL by NRF2** | $\lambda_{X,m_s}$ | 0.67 | $n_{X,m_s}$ | 2 | $X^0_{m_s}$ | 1000 |
| **Inhibition on NRF2 by E-cadherin** | $\lambda_{E,X}$ | 0.33 | $n_{E,X}$ | 2 | $E^0_X$ | 250 |
| **Inhibition on KEAP1 by miR-200** | $\lambda_{\mu_{200},K_s}$ | 0.1 | $n_{\mu_{200},K}$ | 2 | ${\mu_{200}}^0_K$ | 5 |
| **Inhibition on NRF2 by KEAP1** | $\lambda_{K,X}$ | 0.33 | $n_{K,X}$ | 2 | $n_{K,X}$ | 250 |
| **Inhibition on E-cadherin by ZEB** | $\lambda_{Z,E}$ | 0.1 | $n_{Z,E}$ | 2 | $n_{Z,E}$ | 100 |
| **Inhibition on ZEB by E-cadherin** | $\lambda_{E,m_z}$ | 0.8 | $n_{E,m_z}$ | 2 | $n_{E,m_Z}$ | 80 |

**Table SI 2. List of parameters for function $Y$ and $L$.**

| **n (# of miRNA binding sites)** | **0** | **1** | **2** | **3** | **4** | **5** | **6** |
|---|---|---|---|---|---|---|---|
| $l_i$**(hour$^{-1}$)** | 1 | 0.6 | 0.3 | 0.1 | 0.05 | 0.05 | 0.05 |

| | | | | | | | |
|---|---|---|---|---|---|---|---|
| $\gamma_{mi}$(hour$^{-1}$) | 0 | 0.04 | 0.2 | 1 | 1 | 1 | 1 |
| $\gamma_{\mu i}$(hour$^{-1}$) | 0 | 0.005 | 0.05 | 0.5 | 0.5 | 0.5 | 0.5 |
| $n_{\mu_{200}}$ | 6 | | | $n_{\mu_{34}}$ | | | 2 |
| $\mu_{200}^{0}$ | 10K | | | $\mu_{34}^{0}$ | | | 10K |

**Table SI 3. List of other parameters used in EMT model.**

| Synthesis rate | Value (10^3molecules/hour) | Degradation rate | Value (hour$^{-1}$) | Production rate | Value (hour$^{-1}$) |
|---|---|---|---|---|---|
| $g_{\mu_{200}}$ | 2.1 | $\gamma_{\mu_{200}}$ | 0.05 | $k_k$ | 50 |
| $g_{\mu_{34}}$ | 1.35 | $\gamma_z$ | 0.1 | $k_E$ | 50 |
| $g_z$ | 0.1 | $\gamma_{\mu_{34}}$ | 0.05 | $k_X$ | 50 |
| $g_s$ | 0.1 | $\gamma_S$ | 0.125 | | |
| | | $\gamma_K$ | 0.1 | | |
| | | $\gamma_X$ | 0.1 | | |
| | | $\gamma_E$ | 0.1 | | |

#### 4. More results about epigenetic feedback

#### Epigenetic feedback on the inhibition of NRF2 by KEAP1

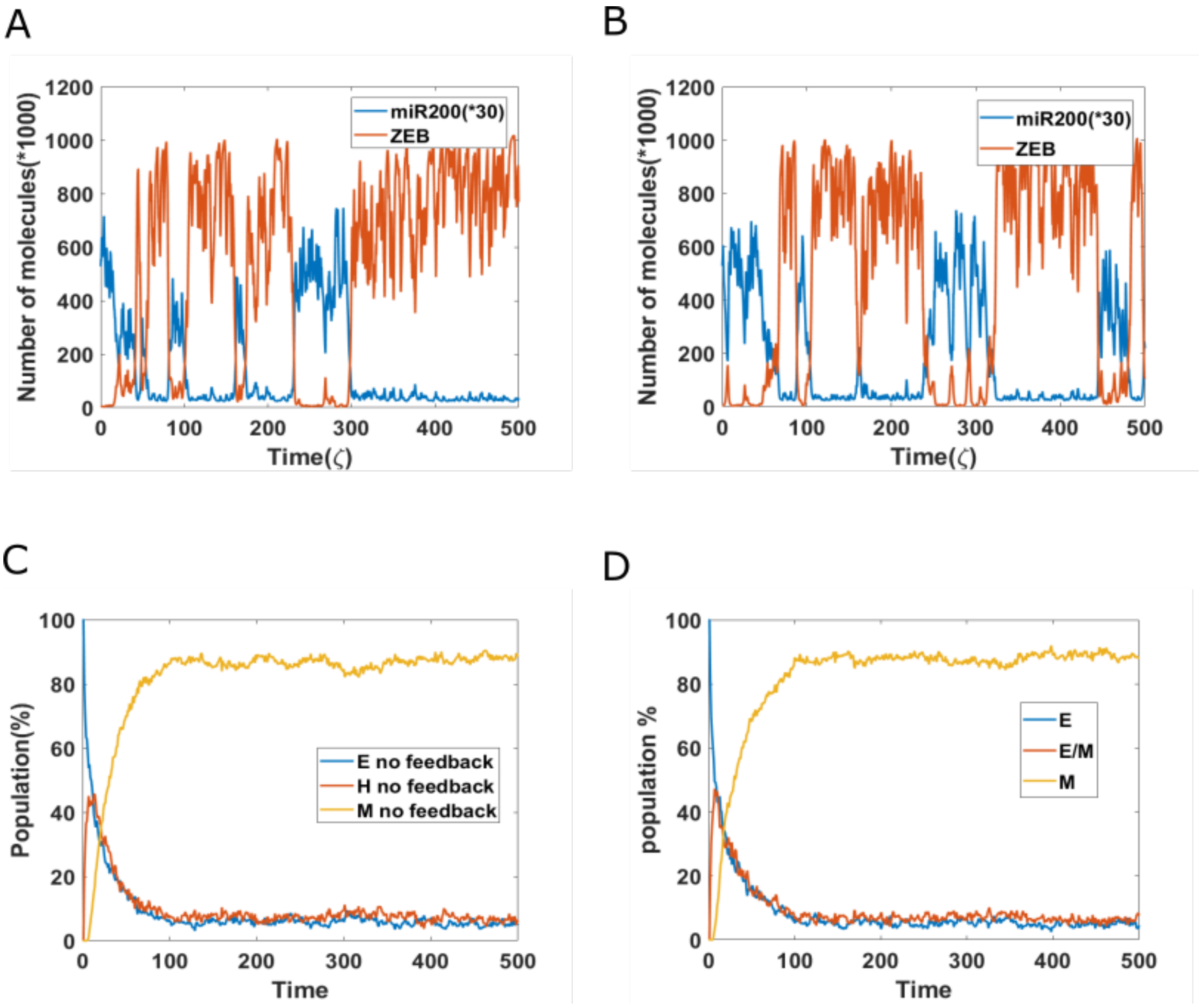


***Figure S1: Epigenetic feedback on inhibition of NRF2 by KEAP1.*** *(A) A sample dynamic plot without epigenetic feedback. (B) A sample dynamic plot with feedback on the inhibition of NRF2 by KEAP1. (C) Simulations showing the population change as a function of time without epigenetic regulation. (D) Same as (C) but now including epigenetic feedback on the inhibitory link from KEAP1 to NRF2.*

**Epigenetic feedback on the inhibition of NRF2 by E-cadherin**

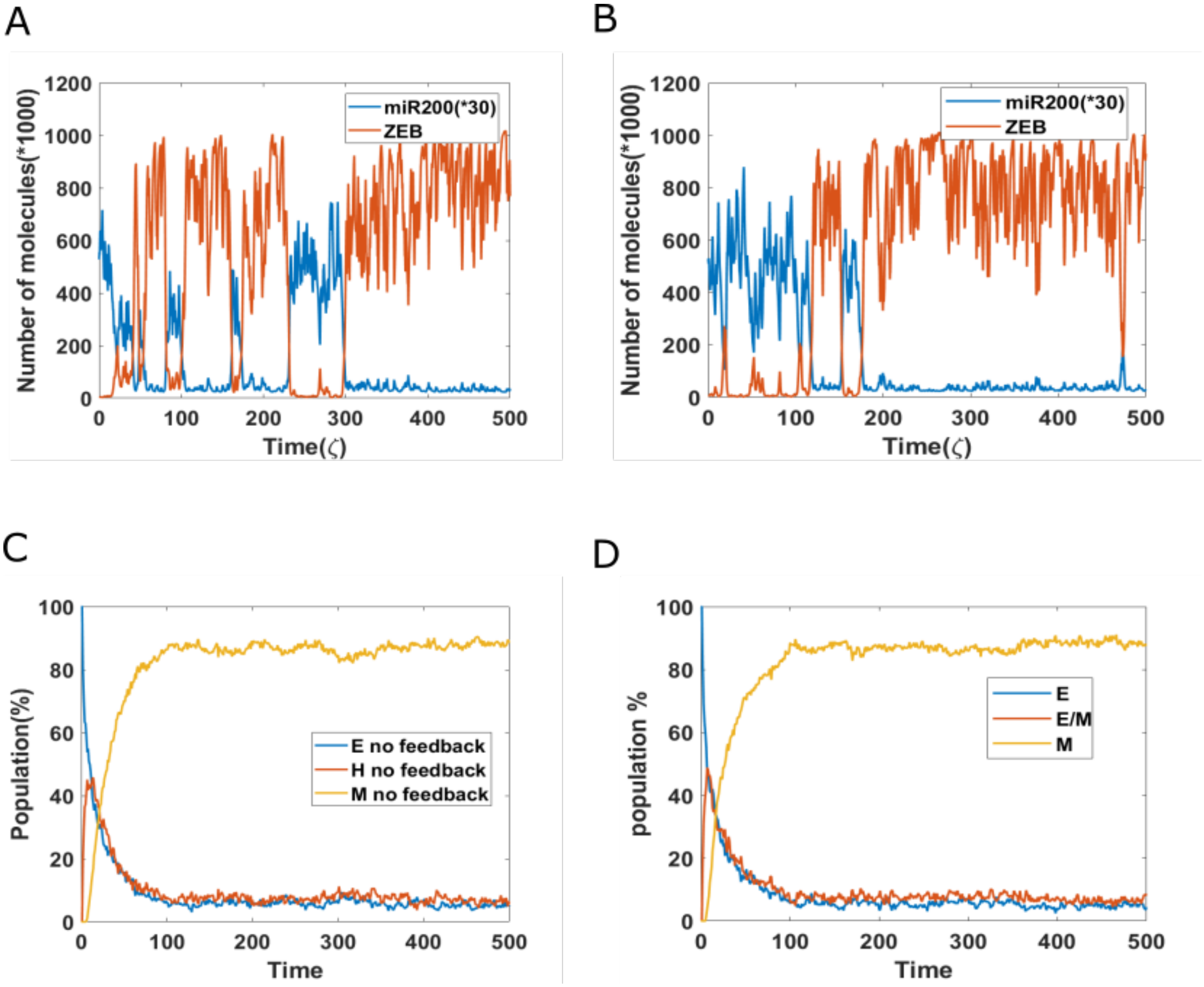


***Figure S2: Epigenetic feedback on inhibition of NRF2 by E-cadherin.*** *(A) A sample dynamic plot without epigenetic feedback. (B) A sample dynamic plot with feedback on the inhibition of NRF2 by E-cadherin. (C) Simulations showing the population change as a function of time without epigenetic regulation. (D) Same as (C) but now including epigenetic feedback on the inhibitory link from E-cadherin to NRF2.*
